# A Cautionary Tail: Changes in Integrin Behavior with Labeling

**DOI:** 10.1101/158618

**Authors:** Catherine G. Galbraith, Michael W. Davidson, James A. Galbraith

## Abstract

Genetic expression of fluorescently labeled proteins is essential to visualizing dynamic behavior within live cells. Recent advances in microscopy have increased resolution to the level where it is now possible to capture individual molecules interacting. However, the criteria for determining whether a fluorescent label perturbs protein function have not undergone a corresponding increase in resolution. The effects of protein labeling on cell function are still judged by whether populations of protein localize and interact with known binding partners. Here we use integrins, bidirectional signal adhesion molecules that regulate interactions between the extracellular matrix and the cytoskeleton through a well-defined series of conformational changes to show that not all labeling strategies are the same. We found that labeling the beta subunit decreased the mobility of individual integrin molecules and the protrusive activity of the entire cell. While integrins with labeled alpha subunits behaved similarly to unlabeled integrins, labeling the beta subunit increased the size of adhesions by elevating integrin affinity and exposing the ligand induced binding domain to change the molecule conformation. Thus, our single molecule and cellular data indicate that the ability of labeled proteins to localize and interact with known binding partners does not guarantee it does not alter protein function. We propose that the behaviors of individual molecules rather than the ensemble behavior of populations need to be considered as criteria to determine if a probe is non-perturbative.

## INTRODUCTION

Genetic expression of fluorescently labeled proteins is essential for visualizing dynamic behavior within live cells. However, it is difficult to know whether labeling has an effect on cell function unless the label induces changes that are significantly larger than inherent cell-to-cell variability. Indeed, labeling is typically assumed to be non-perturbative if the proteins properly localize, respond to known stimulus, and interact with other proteins. Here we present data that challenges that assumption -- molecular labeling strategies that do not inhibit proper localization or interactions, but do induce conformational changes that alter protein affinity for ligand.

We chose integrins as our model because integrins are bidirectional signaling molecules that mediate interactions with the matrix outside of the cell and with the cytoskeleton and signaling molecules inside of the cell by a well-defined series of conformational changes.

Integrins are alpha-beta heterodimers; both subunits are type I transmembrane glycoproteins with large extracellular domains, single spanning transmembrane domains, and short cytoplasmic domains. The extracellular domain is a large globular N-terminal binding head domain, while the transmembrane and cytoplasmic domains are legs or stalks that are severely bent at the knee in the inactive conformation and fully extended in the active conformation when bound to ligand^1,2^. As integrins transition from the inactive to active conformation, they become primed, which is a elevated affinity state where the receptor is not yet bound to ligand^3^. The transition to an elevated affinity state begins with separation of the cytoplasmic and transmembrane domains of the two subunits. As a direct consequence, the interface between the subunits in the tailpiece destabilizes, facilitating straightening of the legs^4^. Mn^2+^ treatment induces extension of the integrin, but results in a mixture of open and closed headpieces, suggesting elevated affinity but incomplete activation. In contrast, ligand binding exclusively produces extended legs with open headpieces, suggesting complete activation^1,2^. These well-defined connections between conformational transitions and molecular behavior provide a well-defined system for studying the effects of labeling on molecular behaviors.

Extensive live-cell studies of GFP labeled integrins have shown that irrespective of whether the label is placed on the alpha or the beta subunit, integrins express on the cell surface and interact with their extracellular and intracellular binding partners. While these studies suggest that integrins are functional^5^, they have been limited to ensemble measures of populations, or measures of individual molecules sequestered into adhesion complexes^6^. We have recently shown that populations of individual unsequestered molecules reveal reproducible molecular behaviors that are obscured by ensemble measures. Therefore, we applied this approach to test whether labeling strategies altered molecular behaviors^7^. To our surprise, we found that molecular mobility as well as the affinity of integrins for matrix changed with labeling strategy. Our results suggest that localization and the ability to interact with binding partners are not adequate metrics to confirm that a label does not perturb protein function – understanding the effects of molecular labeling requires measuring behaviors of individual molecules.

## RESULTS AND DISCUSSION

### Single-Molecule Mobility of Integrins Depends on which Subunit is Fluorescently Labeled

To determine whether labeling one subunit versus the other changes single molecule behavior we expressed both subunits of alpha V beta 3 integrins in CHO-K1 cells, which do not endogenously express either subunit^8^. We left one subunit unlabeled and labeled the other subunit with mEos2, a photoactivatible fluorophore. We then stochastically photo-converted the labeled subunits and used super-resolution microscopy to localize the position of the integrins in live cells from images collected at a frame rate of 40Hz (Supplemental movies S1 and S2). Photo-converted integrins were tracked so that we could calculate the diffusion coefficient and classify molecular mobility behavior as confined, free, or directed diffusion^9,10^. We discovered that for all cells analyzed, the mobility of freely diffusing integrins was statistically lower when the expressed integrins had labeled beta subunits rather than labeled alpha subunits (Fig 1a, b). Integrins confined within adhesion complexes had similar mobilities regardless of which subunit was labeled (Fig 1b, inset). These data indicate that labeling the beta subunit decreases the molecular mobility of the integrins that are not already interacting with either ligand or other proteins.

**Figure 1:**
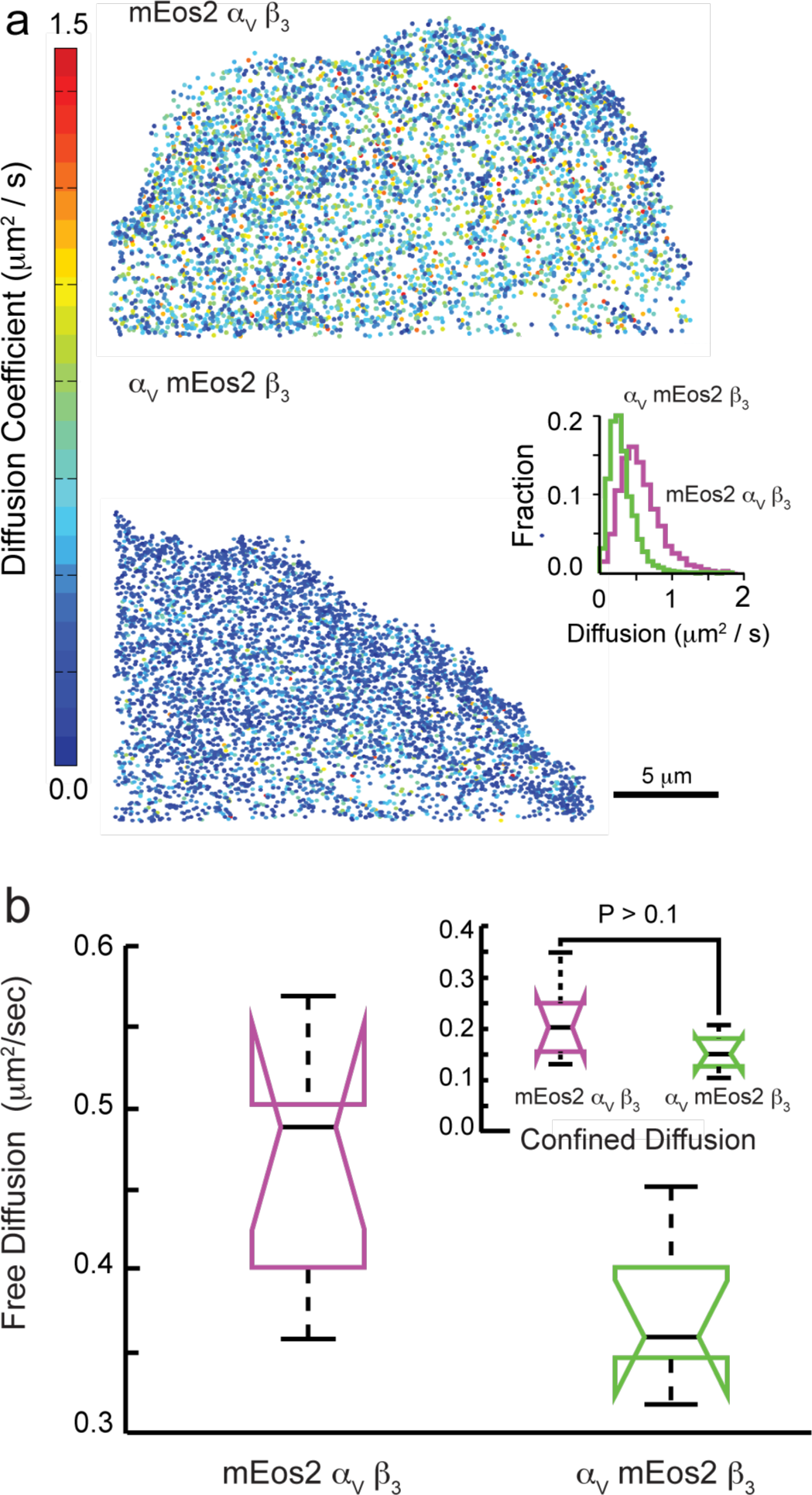
Mobility of Integrins Depends on which Subunit is Fluorescently Labeled. Comparison of mobility of single integrin molecules expressed as either labeled alpha V-unlabeled beta 3 or unlabeled alpha V-labeled beta 3 reveal that labeling the beta subunit slows the mobility of the integrin heterodimer. a) Diffusion coefficient of individual alpha or beta labeled integrin molecules collected over 120 s were color-coded and plotted as points whose centroid indicated the mean location of the integrin. Cells expressing integrins with labeled alpha subunits (top) show populations of integrins have higher mobility (larger diffusion coefficients) compared to populations of integrins with labeled beta subunits (bottom). Inset: Histogram of diffusion coefficients for the cells in panel a reveal that integrins with labeled beta subunits are slower. b) Diffusion coefficients for unconfined movement are significantly slower, p < 0.025, n=5048, 4164 molecules from N = 6 cells for alpha labeled and beta labeled subunits, respectively. Inset: Diffusion coefficients for integrins showing confined movement (integrins within adhesions) are not statistically different, p > 0.1, n=1844, 1323 molecules from N = 6 cells for alpha labeled and beta labeled subunits, respectively.

### Protrusive Activity Changes When the Beta Subunit is Labeled

We then sought to determine if these decreases in integrin mobility that occurred when the beta subunit was labeled also changed dynamic cell behavior when the integrins were labeled with a non-photoactivatible, conventional fluorophore. We replaced the mEos2 labels with mEmerald and 24 hrs post-transfection collected time-lapse images of cells plated on fibronectin. We found that the leading edges of cells transfected with the labeled alpha subunit were consistently more dynamic than the leading edges of cells transfected with the labeled beta subunit (Fig 2a). The quiescence of the leading edge of the cells transfected with the beta subunit labeled is consistent with the notion that labeling the beta subunit increases the affinity of integrins for ligand, and it is also consistent with the more organized adhesions present in the cells transfected with labeled beta subunits (Fig 2b).

**Figure 2:**
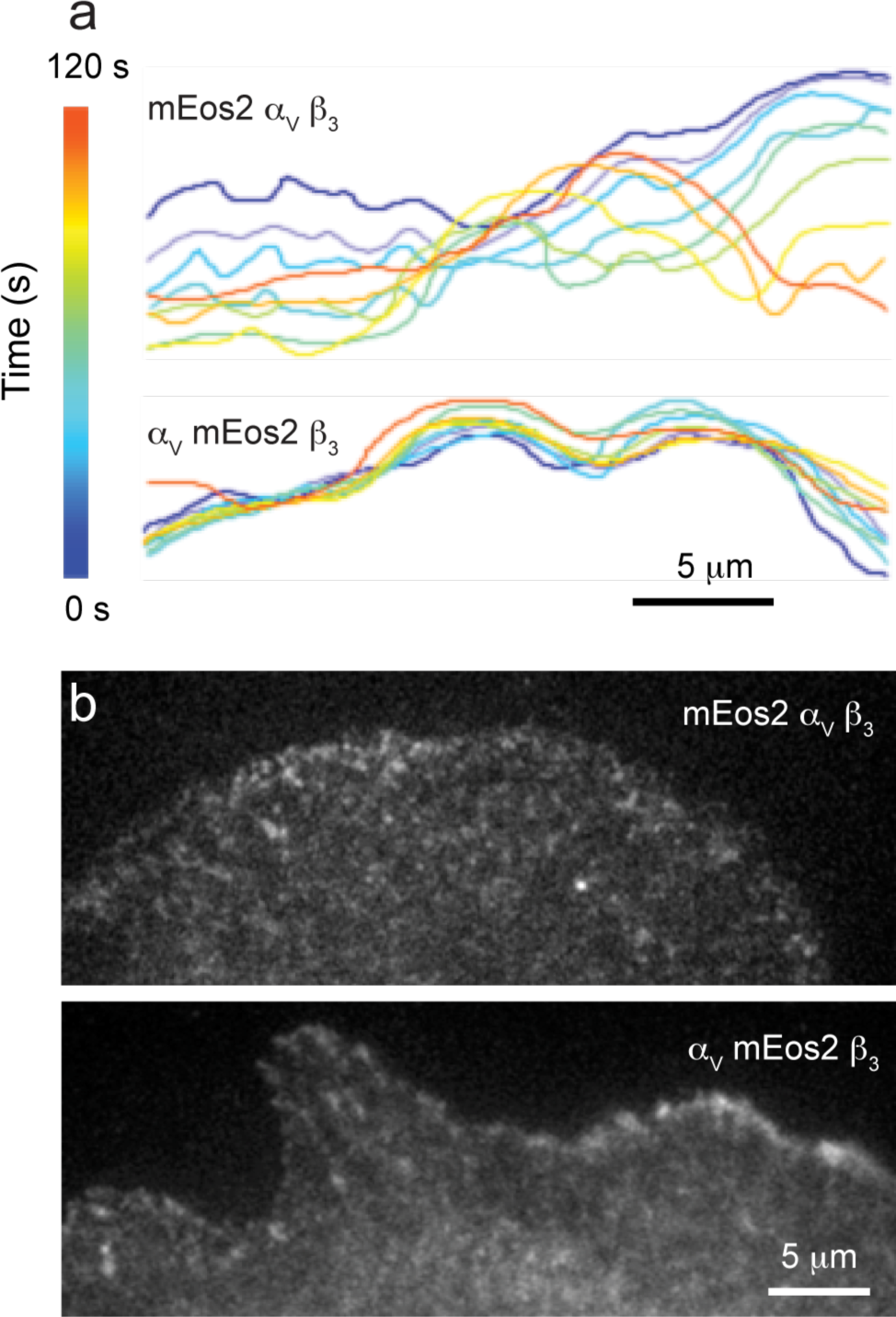
Protrusive Activity Changes When the Beta Subunit is Labeled. Cells expressing integrins with labeled alpha subunits have more protrusive activity and less organized adhesions than cells with labeled beta subunits. a) Representative cell edge contours plotted every 15 s and color coded for time show much more protrusive activity when cells express integrins with labeled alpha subunits in comparison with cells that express integrins with labeled beta subunits. b) Adhesions are smaller when cells express labeled alpha subunits (top) and larger and more organized at the leading edge when cells express labeled beta subunits (bottom).

### Adhesion Size Increases When the Beta Subunit is Labeled

We next investigated whether labeling the beta subunit would alter the size of adhesion complexes, macromolecular scaffolds that connect the cell to matrix and the cytoskeleton via linkages constructed by activated integrins^11^. In addition to transfecting cells with either labeled alpha or beta subunits, we also transfected cells with both subunits unlabeled. We then treated a separate group of the unlabeled integrins with 0.5 mM Mn^2+^ to activate the integrins^11^. Cells transfected with unlabeled integrin were labeled with the human alpha V beta 3 specific LM609 antibody and Alexa 488. All cells were plated on fibronectin-coated coverslips for 3-5 hrs then fixed. We discovered that adhesion size was not statistically different if cells were transfected with unlabeled subunits or with labeled alpha V (Fig 3a). However, adhesions were consistently larger when cells were transfected with labeled beta 3 or when cells expressing unlabeled integrins were treated with Mn^2+^. This data, as well as the similarity in size between the beta labeled integrins and the unlabeled integrins treated with Mn^2+^, further support the interpretation that labeling the beta subunit is activating the integrin.

**Figure 3:**
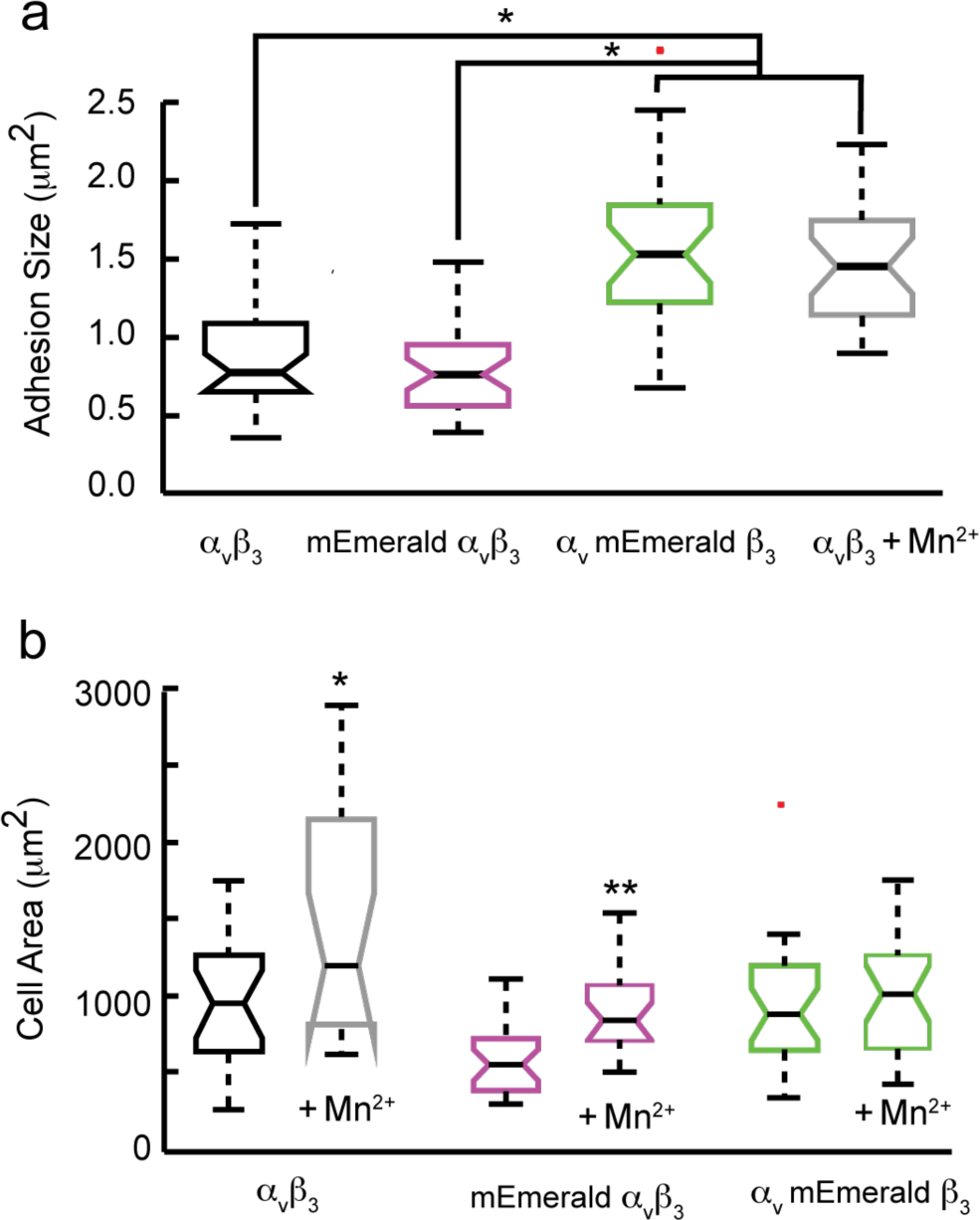
Adhesion Size and Cell Spreading Increases When the Beta Subunit is Labeled. a) Focal adhesion size is larger in cells expressing integrins with labeled beta subunits and in cells expressing integrins with unlabeled subunits that have been treated with Mn^2+^, p<0.0001*. Cells expressing integrins with no labels or labeled alpha subunits have similar sized focal adhesions, p > 0.38, n=33, 35 adhesions from N=6 and 5 cells for unlabeled and alpha labeled subunits, respectively. Cells expressing integrins with the beta subunit labeled or both subunits unlabeled but treated with the integrin activator, Mn^2+^, have similar sized focal adhesions, p > 0.35, n=27 adhesions from N=5 and 4 cells for beta labeled and unlabeled subunits treated with Mn^2+^, respectively. b) Cell spreading increases with integrin activation or with expression of integrins with labeled beta subunits. Cells expressing integrins with various combinations of labeled and unlabeled integrins were allowed to spread for 90 min with and without Mn^2+^ treatment. Mn^2+^ yielded the expected increase in cell spreading (unlabeled: p<0.03*, labeled-alpha: p<0.0005**), except in cells that were expressing integrins with labeled beta subunits. Cells expressing labeled beta subunits did not increase their area in response to Mn^2+^ treatment (p>0.5). N=20, 18, 27, 21, 24, 24 cells for unlabeled, unlabeled Mn^2+^, labeled-alpha, labeled-alpha Mn^2+,^ labeled beta, and labeled-beta Mn^2+^, respectively.

### Whole Cell Response to Matrix Ligand Increases when Beta Subunit is Labeled

We also analyzed the effect of subunit labeling on the interaction of the whole cell with matrix by measuring the ability of cells to spread on matrix-coated surfaces. Cells were again transfected with unlabeled subunits, labeled alpha V, or labeled beta 3 integrin. Approximately 24 hrs post transfection, cells were trypsinized and separated into two groups. One group was treated with Mn^2+^, and both groups were allowed to spread on fibronectin for 30, 60, or 90 min prior to fixation. Treatment of the cells expressing unlabeled integrins or labeled alpha V with Mn^2+^ increased cell spreading, indicating, as expected, that Mn^2+^ activated these integrins. In contrast, treating cells expressing labeled beta 3 integrin with Mn^2+^ did not produce any additional increase cell spreading (Fig 3b). Together these results indicate that labeling the alpha subunit does not activate the integrins, but labeling the beta integrin activates the integrin to a level that is similar to Mn^2+^ treatment.

### Labeling the Beta Subunit Exposes the Ligand Induced Binding Site

To directly test whether labeling increases integrin affinity for ECM, we expressed either unlabeled, alpha subunit labeled, or beta subunit labeled of another integrin, *α*_5_*β*_1_, in CHO-B2 cells, which lack endogenous alpha 5 integrin^12^. We then measured integrin affinity for ligand by quantifying the intensity of 9EG7, a commercially available β1 integrin antibody that detects the ligand-induced binding site (LIBS), which is exposed by Mn^2^ treatment. Integrin affinity was not significantly different when both subunits were unlabeled or when only the alpha subunit was labeled. However, integrins with labeled beta subunits and unlabeled integrins treated with Mn^2+^ both had elevated affinity levels and were not statistically different from each other (Fig 4a). These data demonstrate that labeling the beta subunit elevates the integrin affinity state and induces the same conformational changes as Mn^2+^.

**Figure 4:**
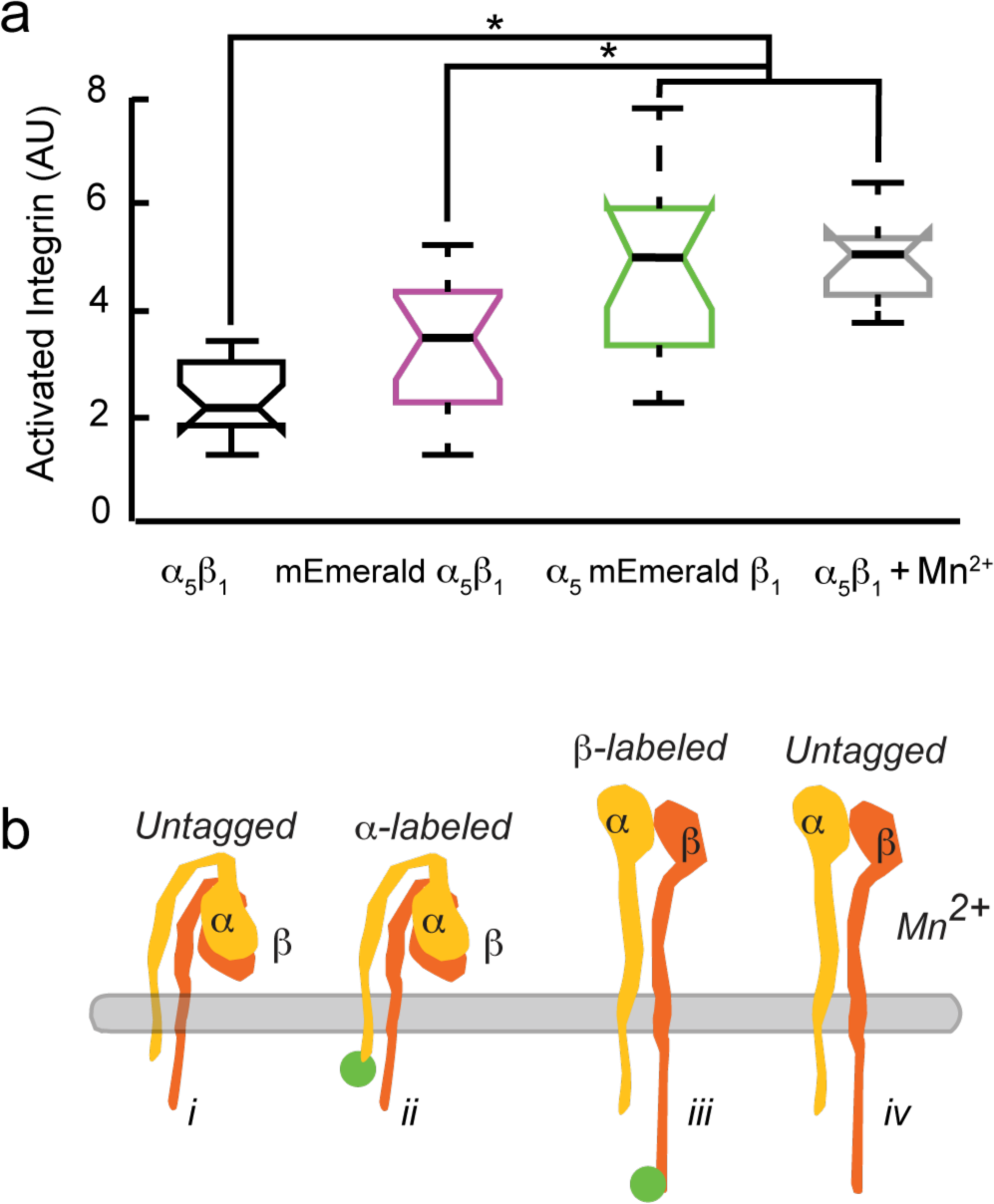
Labeling the Cytoplasmic Tail of the Beta Subunit Increases the Affinity State of Integrin Heterodimers. a) CHO-B2 cells expressing alpha 5 beta 1 integrins were labeled with 9EG7, an antibody that detects the conformational change that occurs when integrins are primed or activated and their affinity for ligand increases. Quantitative immunofluorescence detects a similar conformational change in CHO-B2 cells expressing labeled beta subunits or unlabeled subunits treated with Mn^2+^, but not in untreated cells expressing unlabeled or labeled alpha subunits. N=18, 17, 17, and 13 cells for unlabeled, alpha labeled, beta labeled, and unlabeled and treated with Mn^2+^, respectively. b) Cartoon illustrating the four-different integrin states: i) unlabeled integrin, ii) labeled alpha subunit, iii) labeled beta subunit, and iv) unlabeled integrins exposed to Mn^2+^.

Integrin conformational state is indicative of its ability to bind ligand and organize into adhesions. We found that integrins assembled from unlabeled subunits, labeled alpha subunits, or labeled beta subunits all properly localized to adhesion complexes, indicating that they are all functional. However, measuring the mobility of individual integrin molecules revealed that integrins with labeled beta subunits had a significantly lower diffusion coefficient than integrins with labeled alpha subunits outside of adhesion complexes. These changes at the single molecule level were reflected at the cellular level as less dynamic leading edges, larger adhesions, and larger surface areas in spreading assays compared to cells expressing labeled alpha subunits. These changes are functionally and statistically significant, but were not so morphologically abnormal that they would be identified as outside of normal cell variability. However, by comparing these cells to cells expressing unlabeled integrins treated with Mn^2+^, our data suggested that labeling the beta subunit was activating the integrin (Fig 4b).

An earlier study noted that CHO-K1 cells expressing GFP-labeled alpha IIb (beta 3) or alpha IIb (GFP-labeled beta 3) were more likely to spontaneously aggregate in the presence of soluble fibrinogen, the ligand for alpha II (beta 3) integrin^5^. However, that study also noted that both labeling strategies did not inhibit the formation of adhesions and cell spreading. Therefore, they concluded that labeling either subunit was non-perturbative^5^. Our single molecule results, as well as our comparison of focal adhesion size and cell spreading in untreated cells and in cells with integrins activated by exposure to Mn^2+^ all suggest that labeling the beta subunit changes the function of the expressed integrin by increasing its affinity for ligand.

Further evidence for the interpretation that labeling the beta subunit increases integrin affinity for ligand comes from quantification of the exposure of the LIBS domain in untreated cells and in cells exposed to Mn^2+^. Here our data suggests that labeling the beta subunit induced separation of the alpha and beta cytoplasmic tails to expose the LIBS domain, producing a response similar to Mn^2+^ treatment. These results point to a cautionary tale -- localization and the ability to interact with binding partners are not adequate to confirm that a label is truly non-perturbative. Our data suggest that functionality of molecules needs to be evaluated by measuring molecular behaviors and that localization and reorganization on the cellular level are insufficient metrics for guaranteeing that labeling does not alter molecular functionality.

## MATERIALS AND METHODS

### Cell culture and transfection

CHO-K1 cells, which do not express endogenous alpha V or beta 3 integrins (ATCC) and CHO-B2 cells, which do not express endogenous alpha 5 were grown in DMEM-F12 supplemented with 10% FBS. Cells were transfected with either human alpha V and beta 3 (CHO-K1) or alpha 5 and beta 1 (CHO-B2) using a Nucleofector II (Lonza) and Ingenio (Mirus) transfection reagents following manufacturer’s protocols. Unlabeled integrins were in either pcDNA3.1 vectors (alpha V, beta 3, and alpha 5) or pRK5 (beta 1), and labeled vectors (mEos2 or Emerald) were constructed as previously described^7^. The labeled vectors are available through Addgene as mEos2-Alpha-V-integrin-N-25, mEmerald-Alpha-V-integrin-N-25, mEos2-Integrin-Beta3-N-18, mEmerald-Beta3-N-18, mEmerald-Alpha5-Integrin-12, and mEmerald-Beta1-N-18. The unlabeled alpha V and beta 3 subunits were gifts from Mark Ginsberg (UCSD). Cells transfected with mEos2 labeled subunits were also transfected with an EFGP vector to identify edge contours in live cell experiments. Cells were plated on plasma-etched cover glass that had been silanized and coated overnight with either 5µg/ml human plasma fibronectin (CHO-K1) or 10µg/ml human plasma fibronectin (CHO-B2).

### Cell Spreading and Integrin Activation Assays

Approximately 24 hrs after transfection, cells were trypsinized and plated for 30, 60, or 90 min for adhesion assays or 4 hrs for spreading assays. Cells were then fixed with 2% paraformaldehyde in PHEM^13^. Cells that were transfected with unlabeled integrins were co-transfected with an empty Emerald vector for visualization of the cell perimeter. For activation, treatment with 0.05mM Mn^2+^ was initiated 5 min prior to plating and was maintained throughout the spreading assay^14^. To detect adhesions in cells transfected with unlabeled integrins, cells were labeled with LM609 (Millipore) prior to secondary labeling with Alexa 488.

To quantify the amount of integrin activation transfected cells were plated overnight prior to fixation and labeling with 9EG7 (BD Pharmigen) prior to secondary labeling with Alexa 647. 9EG7 interacts with high affinity mouse and human beta 1 integrin^3^, binding the ligand-induced binding epitope exposed by treatment with Mn^2+15^, 9EG7 does not interact with endogenous hamster integrin. Activation with 0.05mM Mn^2+^ was initiated 1 hr prior to fixation.

### Microscopy

All imaging experiments were performed on an Olympus IX71 with a 60X 1.49 NA objective using TIRF illumination. To create the TIRF beam four laser lines (405, 488, 561, 633 nm) (Coherent) were merged and introduced through free space into the TIRF illumination port of the microscope. Position of the beam in the back aperture of the objective was motorized to ensure repeatability of the penetration depth of the evanescent TIRF wave. For the single molecule experiments a subpopulation of the mEos2 labeled molecules was stochastically excited with a low level of 405 nm activation and 561 nm excitation light (5 µW and 2.5 mW at the back aperture respectively). In these experiments cell edges were identified by collection of an image of unconjugated EGFP. In the case of live cell single-molecule experiments, every 10 sec (400 frames), the excitation light was switched to 488 nm (100 µW at the back aperture) using an acousto-optic tunable filter (AOTF, AA Opto-Electronic). Live cell experiments were imaged at 37° C for a minimum of 5 min, and cells did not display any abnormal morphology or decreased motility at the end of this interval. Images were acquired at a final magnification of 111 nm/pixel with an Andor 897 EMCCD camera using an exposure time of 25 ms. For the activation quantification experiments, the laser power was maintained constant for all images.

### Image processing, single molecule analysis, and statistics

The cell edge and adhesions were detected by thresholding images in Fiji^16^ after smoothing with a 1 pixel Gaussian kernel sigma to reduce noise. Canny edge detection was used on the whole cell images after thresholding to obtain cell contours.

Single molecule analysis was performed using uTrack software^17^ to localize and track individual mEos2 integrin molecules. Only molecules localized to better than 25nm precision were used for mobility analysis. Diffusion coefficients for tracks greater than 20 frames were analyzed as previously described and classified as either confined, freely diffusing or undergoing directed movement (i.e. drift)^7,10^. An average of between 6000 and 7000 molecules per cell was analyzed with a minimum of 6 cells per experimental group.

One-way ANOVA analysis was performed on all experimental groups. Sheffer post-hoc multiple comparison test was used to identify which treatments significantly differed from each other.

The datasets generated and analyzed during the current study are available from the corresponding author on reasonable request.

## AUTHOR CONTRIBUTIONS

C.G.G. and J. A.G. designed experiments. C.G.G. performed experiments and J.A.G. performed analysis. M. W. D. designed and created the labeled constructs. All authors discussed results. C.G.G. and J.A.G. wrote the manuscript.

## REFERENCES

1 Takagi, J., DeBottis, D. P., Erickson, H. P. & Springer, T. A. The role of the specificity-determining loop of the integrin beta subunit I-like domain in autonomous expression, association with the alpha subunit, and ligand binding. Biochemistry 41, 4339–4347 (2002).

2 Takagi, J., Petre, B. M., Walz, T. & Springer, T. A. Global conformational rearrangements in integrin extracellular domains in outside-in and inside-out signaling. Cell 110, 599–511 (2002).

3 Galbraith, C. G., Yamada, K. M. & Galbraith, J. A. Polymerizing actin fibers position integrins primed to probe for adhesion sites. Science 315, 992–995, doi:10.1126/science.1137904 (2007).

4 Carman, C. V. & Springer, T. A. Integrin avidity regulation: are changes in affinity and conformation underemphasized? Curr Opin Cell Biol 15, 547–556 (2003).

5 Plancon, S., Morel-Kopp, M. C., Schaffner-Reckinger, E., Chen, P. & Kieffer, N. Green fluorescent protein (GFP) tagged to the cytoplasmic tail of alphaIIb or beta3 allows the expression of a fully functional integrin alphaIIb(beta3): effect of beta3GFP on alphaIIb(beta3) ligand binding. Biochem J 357, 529–536 (2001).

6 Rossier, O. et al. Integrins beta1 and beta3 exhibit distinct dynamic nanoscale organizations inside focal adhesions. Nat Cell Biol 14, 1057–1067, doi:10.1038/ncb2588 (2012).

7 Jaqaman, K., Galbraith, J. A., Davidson, M. W. & Galbraith, C. G. Changes in single-molecule integrin dynamics linked to local cellular behavior. Molecular biology of the cell 27, 1561–1569, doi:10.1091/mbc.E16-01-0018 (2016).

8 Xu, X. et al. The genomic sequence of the Chinese hamster ovary (CHO)-K1 cell line. Nat Biotechnol 29, 735–741, doi:10.1038/nbt.1932 (2011).

9 Jaqaman, K. et al. Cytoskeletal control of CD36 diffusion promotes its receptor and signaling function. Cell 146, 593–606, doi:10.1016/j.cell.2011.06.049 (2011).

10 Ewers, H. et al. Single-particle tracking of murine polyoma virus-like particles on live cells and artificial membranes. Proc Natl Acad Sci U S A 102, 15110–15115, doi:10.1073/pnas.0504407102 (2005).

11 Ballestrem, C., Hinz, B., Imhof, B. A. & Wehrle-Haller, B. Marching at the front and dragging behind: differential alphaVbeta3-integrin turnover regulates focal adhesion behavior. J Cell Biol 155, 1319–1332, doi:10.1083/jcb.200107107 (2001).

12 Wu, C., Bauer, J. S., Juliano, R. L. & McDonald, J. A. The alpha 5 beta 1 integrin fibronectin receptor, but not the alpha 5 cytoplasmic domain, functions in an early and essential step in fibronectin matrix assembly. J Biol Chem 268, 21883–21888 (1993).

13 Galbraith, C. G., Skalak, R. & Chien, S. Shear stress induces spatial reorganization of the endothelial cell cytoskeleton. Cell Motil Cytoskeleton 40, 317–330, doi:10.1002/(SICI)1097-0169(1998)40:4-317::AID-CM1>3.0.CO;2-8 (1998).

14 Humphries, M. J. Cell-substrate adhesion assays. Curr Protoc Cell Biol Chapter 9, Unit 9 1, doi:10.1002/0471143030.cb0901s00 (2001).

15 Bazzoni, G., Shih, D. T., Buck, C. A. & Hemler, M. E. Monoclonal antibody 9EG7 defines a novel beta 1 integrin epitope induced by soluble ligand and manganese, but inhibited by calcium. J Biol Chem 270, 25570–25577 (1995).

16 Schindelin, J. et al. Fiji: an open-source platform for biological-image analysis. Nat Methods 9, 676–682, doi:10.1038/nmeth.2019 (2012).

17 Jaqaman, K. et al. Robust single-particle tracking in live-cell time-lapse sequences. Nat Methods 5, 695–702, doi:10.1038/nmeth.1237 (2008).

